# The PUF RNA-binding protein, FBF-2, maintains stem cells without binding to RNA

**DOI:** 10.1101/2024.10.25.620246

**Authors:** Brian H. Carrick, Sarah L. Crittenden, MaryGrace Linsley, Stephany J. Costa Dos Santos, Marvin Wickens, Judith Kimble

## Abstract

Like all canonical PUF proteins, *C. elegans* FBF-2 binds to specific RNAs via tripartite recognition motifs (TRMs). Here we report that an FBF-2 mutant protein that cannot bind to RNA, is nonetheless biologically active and maintains stem cells. This unexpected result challenges the conventional wisdom that RBPs must bind to RNAs to achieve biological activity. Also unexpectedly, FBF-2 interactions with partner proteins can compensate for loss of RNA-binding. FBF-2 only loses biological activity when its RNA-binding and partner interactions are both defective. These findings highlight the complementary contributions of RNA-binding and protein partner interactions to activity of an RNA-binding protein.

## Introduction

RNA-binding proteins (RBPs) pervade eukaryotic biology, from yeast to humans (Gerstberger et al. 2014). The conventional view is that RNA binding is essential for RBP biological activity (Lunde et al. 2007; Corley et al. 2020). Based on this view, mutations in RNA-binding residues are often mutated to abolish RBP activity (e.g. Siomi et al. 1994; Daigle et al. 2013).

The RNA-binding domain (RBD) of PUF (<u>Pu</u>milio and <u>F</u>BF) RNA-binding proteins is defined by eight “PUF repeats” (Fig. 1A-C). These repeats form a crescent that binds to RNA on its inner surface and binds to protein partners on its outer surface (Fig. 1A-C) (Zamore et al. 1997; Edwards et al. 2001; Wang et al. 2001; Campbell et al. 2012b; Friend et al. 2012; Weidmann et al. 2016; Qiu et al. 2019; Carrick et al. 2024). *In vitro*, RNA-binding of the PUF RBP relies on tripartite recognition motifs (TRMs), which reside in each PUF repeat. TRM motifs recognize and bind to nucleotides within the PUF RNA binding element and hence define PUF sequence specificity (Wang et al. 2002; Campbell et al. 2012a) (Fig. 1B-D). However, their biological role has not yet been explored.

**Figure 1:**
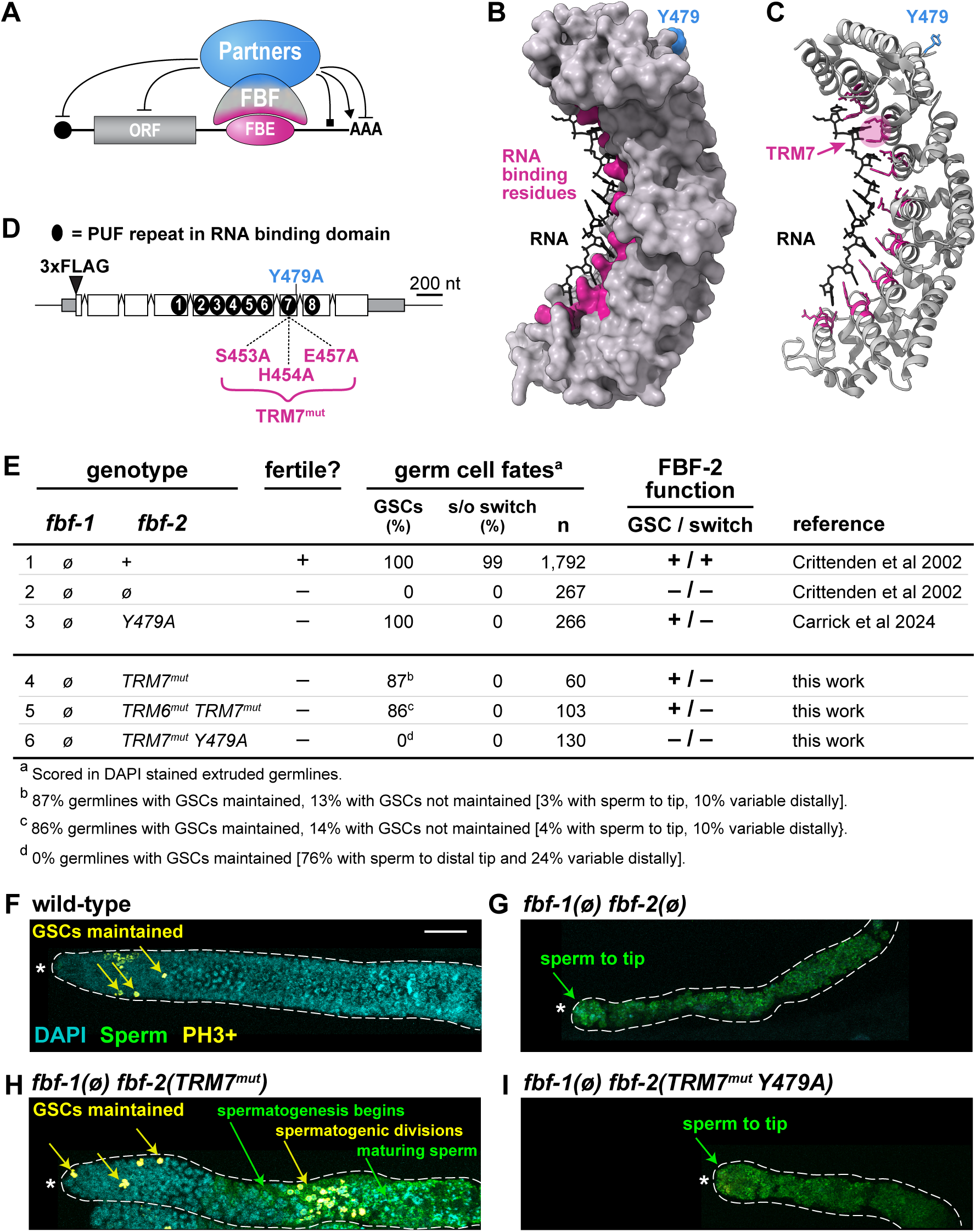
FBF-2 *in vivo* function requires both RNA- and protein-binding residues. (A) FBF binds to RNA and protein partners via distinct interfaces (magenta for RNA, blue for partners) to control various activities: RNA repression, blunted end; RNA activation, arrowhead; RNA binding square end. ORF, open reading frame; FBE, FBF-binding element. (B-C) Crystal structure of FBF-2 (B, surface; C, ribbon) binding to RNA (PDB: 3K5Y) (Wang et al. 2009). RNA binding residues, magenta; Y479 partner interface, blue. Tripartite Recognition Motifs (TRMs) in each PUF repeat mediate RNA-binding; TRM7 is highlighted in C. (D) *fbf-2* mRNA and FBF-2 protein features. Untranslated regions (gray boxes); coding regions (white boxes), introns (peaked lines), PUF repeats (black ovals). Sites for insertion of 3xFLAG and relevant mutations are indicated. (E) FBF-2 TRM mutant effects on germline fates, scored in DAPI-stained extruded gonads. GSCs, % of germlines with stem cells maintained to the distal end; s/o switch, % of germlines with a successful sperm-to-oocyte switch. n, number gonads scored. (F-I) Representative z-projection images of extruded adult gonads, stained for DNA (DAPI, cyan), sperm (αSP56, green), and a cell cycle marker, phosphohistone H3 (αPH3, yellow). αPH3 marks cells in both mitotic and meiotic G2/M phase. GSC maintenance is inferred from mitotic divisions in the distal gonad (αPH3-positive staining, yellow arrows); spermatogenic meiotic divisions occur more proximally (αPH3-positive staining, green arrows) where SP56-positive staining indicates sperm differentiation. Dotted line marks gonad boundary; asterisk marks distal end. 20µm scale bar in (F) applies to (F-I).

In addition to binding RNAs, PUF proteins also interact with proteins that help set the RNA regulatory mode, for example repression (e.g. Ccr4-Not complex) (Goldstrohm et al. 2007; Webster et al. 2019) or activation (e.g. GLD-3, GLD-2) (Suh et al. 2009) and can also affect PUF RNA binding affinity (e.g. CPEB, Nanos, LST-1) (Campbell et al. 2012a; Weidmann et al. 2016; Qiu et al. 2019) (Fig. 1A). One interaction site used by multiple protein partners occurs at the R7/R8 loop between PUF repeats 7 and 8 on the outer surface of the PUF RBD crescent (Fig. 1B-C) (Campbell et al. 2012b; Wu et al. 2013; Qiu et al. 2019; Qiu et al. 2023; Carrick et al. 2024). Within the R7/R8 loop of nematode FBF-2, a paradigmatic PUF protein, a single tyrosine (Y479) stands out as critical for binding multiple partners; indeed, its alanine substitution (Y479A) abolishes partner binding when assayed *in vitro*, in yeast and in nematodes (Campbell et al. 2012b; Carrick et al. 2024).

FBF-2 drives two major biological functions in the *C. elegans* germline: maintenance of germline stem cells (GSCs) and promotion of the sperm-to-oocyte cell fate switch (s/o switch) (Zhang et al. 1997; Crittenden et al. 2002; Lamont et al. 2004). FBF-2 and its nearly identical counterpart FBF-1 are biologically redundant and can substitute for each other to accomplish these two functions. Thus, FBF-2 null mutants are fertile when FBF-1 is wild-type (Fig. 1E, row 1), but sterile when FBF-1 is removed, due to loss of GSCs and failure of the s/o switch (Fig. 1E, row 2) (Crittenden et al. 2002). We previously reported the molecular and biological effects of a partner defective Y479A mutant (Carrick et al. 2024). When FBF-1 was wild-type and germlines essentially normal, the FBF-2(Y479A) mutant protein changed the RNA binding, as assayed by eCLIP. That shift revealed that partner interactions modulate FBF-2 RNA-binding strength at specific FBF binding elements (FBEs). On the other hand, when FBF-1 was gone, Y479A was able to maintain stem cells but failed to promote the s/o switch (Fig. 1E, row 3) (Carrick et al. 2024). Therefore, Y479-dependent partnerships are essential for one FBF-2 biological activity but not both.

Here, we tested whether the Y479A mutant protein must bind RNA to accomplish its role in GSC maintenance. Unexpectedly, we found that loss of RNA binding on its own does not abolish FBF-2 activity and that Y479-dependent partnerships compensate for loss of RNA binding. More broadly, our findings raise the possibility that other RBPs lacking the ability to bind RNA may nonetheless retain biological activity.

## Results and Discussion

### FBF-2 RNA-binding mutants retain biological function

We predicted that the FBF-2 mutant, Y479A, would rely on RNA-binding for its ability to maintain stem cells. To test that prediction, we mutated three key TRM residues in PUF-repeat 7 (S453A H454A E457A, Fig. 1D) of an endogenous, FLAG-tagged, *fbf-2* gene, thus creating the TRM7^mut^ mutant. We chose these residues because any one of the three TRM7 alanine substitutions abolished FBF-2 RNA binding, when assayed *in vitro* or by yeast three-hybrid (Valley et al. 2012); moreover, alanine substitutions of these same TRM residues in the fly Pumilio protein were used to eliminate RNA binding in reporter assays (e.g. Weidmann and Goldstrohm 2012).

Before making TRM7^mut^ changes in the Y479A mutant, we introduced them into wild-type FBF-2 and scored for its two major biological activities, GSC maintenance and the s/o switch (see Introduction). Our expectation was that loss of RNA-binding would destroy both activities and that TRM7^mut^ would behave like an *fbf-2(ø)* mutant. To score FBF-2 activities, we removed FBF-1 so defects were not masked by redundancy. We examined phenotypes with DAPI (Fig. 1E) and immunostaining (Fig. 1F-I). As expected, wild-type *fbf-2(+)* maintained GSCs and promoted the s/o switch (Fig. 1E row 1, Fig. 1F), while *fbf-2(ø)* lost both functions (Fig. 1E row 2, Fig. 1G). To our surprise, TRM7^mut^ did not behave like the null (Fig. 1E, compare rows 2 and 4; compare Fig. 1G and 1H). Most (87%) TRM7^mut^ adults retained GSCs, demonstrating that contrary to expectation, this mutant retains biological activity. By contrast, all (100%) TRM7^mut^ adults lost the s/o switch (Fig. 1E, line 4; 1H). The TRM7^mut^ was thus able to exert one function (GSC maintenance), but not another (s/o switch), much like Y479A (Fig. 1E compare rows 3 and 4). Regardless, the key conclusion is that TRM7^mut^ retains biological activity – it is sufficient for GSC maintenance. To push the limits of this unexpected result, we mutated the key TRM residues in the neighboring sixth PUF repeat (TRM6: N415A, Y416A, Q419A) to create TRM6^mut^ TRM7^mut^ double mutants. However, TRM6^mut^ did not enhance the TRM7^mut^ phenotype (Fig. 1E compare rows 4 and 5), consistent with TRM7^mut^ being sufficient to abolish RNA-binding (see Fig. 3 for molecular confirmation).

### FBF-2 RNA binding and partner interactions both promote GSC self-renewal

Our original question was whether Y479A requires RNA binding to maintain GSCs. To address that question, we generated TRM7^mut^ Y479A, a double mutant that removes both RNA-binding and Y479A-dependent partner interactions. This double mutant confirmed our expectation. TRM7^mut^ Y479A mimicked an FBF-2 molecular null when assayed without FBF-1 (Fig. 1E, compare rows 2 and 6). Thus, both *fbf-2(ø)* and Y479A TRM7^mut^ mutant germlines lacked GSCs at the distal end when assayed with DAPI. This was confirmed by the absence of staining with a marker for dividing cells (Fig. 1I).

Although both *fbf-2(ø)* and TRM7^mut^ Y479A mutant germlines lacked GSCs, we did find minor differences. All *fbf-2(ø)* and most (76%) Y479A TRM7^mut^ mutant germlines had mature sperm all the way to their distal end (Fig. 1G; Fig. 1I); however, some (24%) Y479A TRM7^mut^ germlines contained cells that had not become mature sperm distally, suggesting marginal activity in some germlines. We speculate this residual activity may be due to partner interactions that occur outside of the R7/R8 loop (e.g. the Ccr4-Not complex or Argonaute). Regardless, no Y479A TRM7^mut^ double mutants were able to maintain GSCs. We conclude that Y479A does indeed rely on RNA binding to maintain GSCs, and that unexpectedly Y479-dependent partner interactions compensate for loss of RNA-binding for that same biological function.

### TRM7^mut^ Y479A behaves like a null when assayed in the presence of FBF-1

We next investigated TRM7^mut^ and TRM7^mut^ Y479A in the presence of wild-type FBF-1, because these germlines were essentially normal, allowing us to test for effects on protein stability and cellular distribution, and because other *fbf-2* mutants had minor defects in this situation (see below). By immunostaining, both the levels and distribution of TRM7^mut^ and TRM7^mut^ Y479A mutant proteins were comparable to wild-type (Fig. 2A, also see Fig. 3A). Therefore, any changes are likely due to an effect on activity rather than stability.

**Figure 2:**
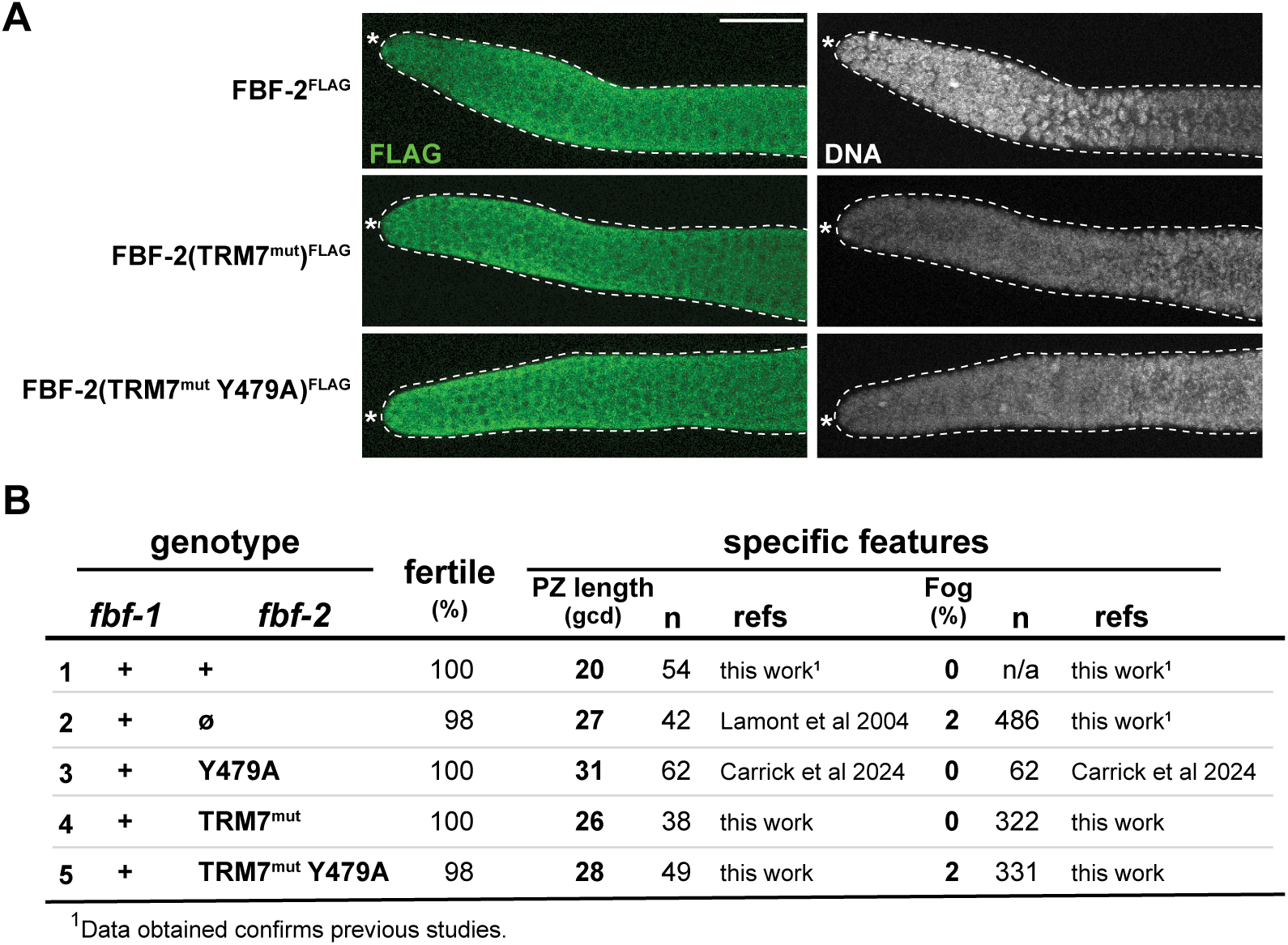
Y479A TRM7^mut^ behaves like a null when assayed in presence of wild-type FBF-1. (A) Representative z-projection images of extruded adult gonads, stained for FLAG:FBF-2 (αFLAG, green) and DNA (DAPI, gray). Dotted line marks gonad boundary; asterisk marks distal end. 20µm scale bar in the top left image applies to all images. (B) FBF-2 mutant defects in the presence of wild-type FBF-1. PZ, progenitor zone size; fertile, animals capable of producing self-progeny. All animals that were not fertile were feminized (only oocytes, Fog).

**Figure 3:**
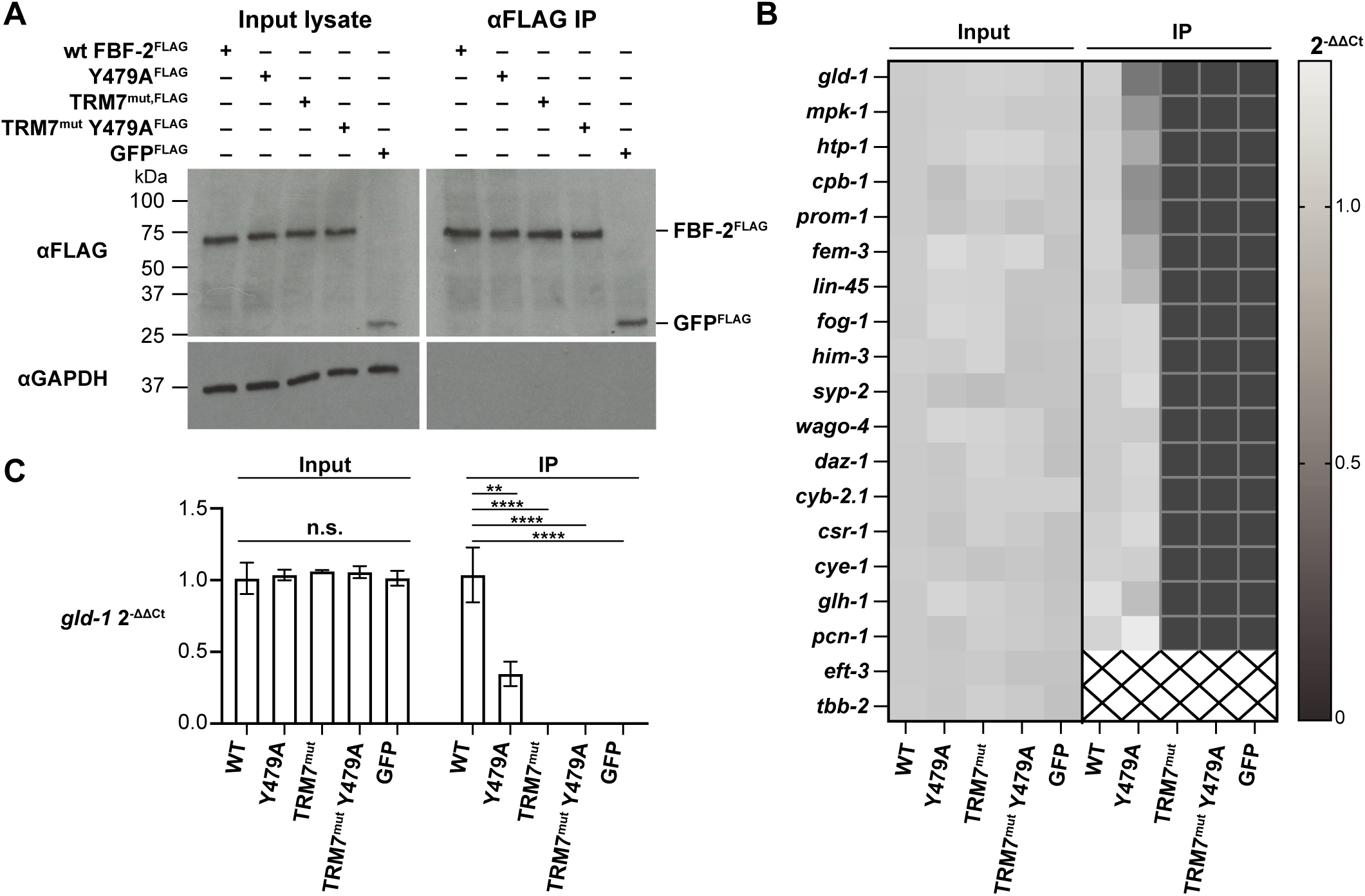
TRM7mut eliminates FBF-2 binding to target RNAs *in vivo*. (A) TRM mutations do not change FBF-2 protein abundance or immunoprecipitation efficiency. Representative western blot for RIP-qPCR experiment. Left, input lysates (1%); right, FLAG IP (1%). FLAG tagged FBF-2 variants and GFP are immunoprecipitated; negative control (GAPDH) is not immunoprecipitated. (B) TRM mutations abolish FBF-2 mRNA binding. Heatmap depicts results from quantitative PCR of FBF target mRNAs and control mRNAs after αFLAG IP, using 3xFLAG::FBF-2 for variants and 3xFLAG::GFP for the control. Mean mRNAs abundance in input (left) and Ips (right) was calculated with the comparative CT method (2-ΔΔCT) (Schmittgen and Livak 2008), using rps-25 for normalization and making all comparisons to the wild-type sample. 2-ΔΔCT = 1, no change in mRNA level compared to wild-type; 2-ΔΔCT < 1, less mRNA than wild-type; 2-ΔΔCT > 1 more mRNA than wild-type (grey level scale indicated to right of heatmap). Because no specific signal was seen for negative controls *eft-3* and *tbb-2* in the wild-type IP sample, 2-ΔΔCT was not calculated (boxes containing black X). (C) Effect of FBF-2 mutations on binding to *gld-1* RNA. Example bar graph of 2-ΔΔCT values for one of the target RNAs tested, *gld-1*. No significant differences in *gld-1* levels were seen in the input. The *gld-1* RNA abundance is significantly different in immunoprecipitated samples. ** p = 0.0017, **** p < 0.0001.

What were the minor defects of *fbf-2* mutants seen previously when assayed in the presence of wild-type FBF-1? First was a lengthening of the Progenitor Zone (PZ), a region in the distal germline that includes GSCs and GSC daughters that have just begun to differentiate. PZ length increased from ∼20 germ cell diameters (gcd) in wild-type animals to >25 gcd in both *fbf-1(+) fbf-2(*ø) and *fbf-1(+) fbf-2(*Y479A) (Fig. 2B, rows 1-3) (Lamont et al. 2004; Carrick et al. 2024). Second was an increase in percentage of feminized germlines (Fog), from zero in wild-type to 2% in *fbf-1(+) fbf-2(*ø) (Fig. 2B, row 1 and 2); this defect was not seen for Y479A (Fig. 2 row 3) (Carrick et al. 2024).

For the TRM7^mut^ single and TRM7^mut^ Y479A double mutants, PZ length increased (Fig. 2B, rows 4,5), much like for FBF-2 null and Y479A mutants (Fig. 2A, row 2 and 3). By contrast, TRM7^mut^ or Y479A single mutants had no feminized Fog germlines (Fig. 2B, rows 4, 3), like wild-type, but TRM7^mut^ Y479A double mutants did generate 2% Fog germlines (Fig. 2B, row 5), like FBF-2 null (Fig. 2B, row 2). We conclude that activity of the TRM7^mut^ Y479A double mutant is comparable to that of an FBF-2 null, both in the presence of FBF-1 (Fig. 2B) and in the absence of FBF-1 (Fig. 1E).

### TRM7^mut^ abrogates RNA binding in vivo

To test whether TRM7^mut^ abolishes RNA binding activity in nematodes, we used RNA immunoprecipitation followed by quantitative PCR (RIP-qPCR) to assess FBF-2 binding to RNA targets *in vivo*. Importantly, these experiments were done using animals with normally organized and functional germlines due to the presence of wild-type FBF-1 (Fig. 2). We performed three biological replicates for each of five FLAG-tagged proteins: wild-type FBF-2, Y479A, TRM7^mut^, TRM7^mut^ Y479A, and GFP as a negative control (Fig. 3A). These FLAG-tagged proteins were expressed at comparable levels and immunoprecipitated with similar efficiency (Fig. 3A). After extracting RNAs that co-immunoprecipitated with the FLAG-tagged proteins and converting them to cDNA, we employed qPCR to probe for 20 different RNAs, including 17 FBF-2 target RNAs and two non-target RNAs (*eft-3* and *tbb-2*). We used another non-target RNA (*rps-25*) for normalization using the comparative C_T_ method (Schmittgen and Livak 2008). Each biological replicate was assessed in technical triplicate, and RNA levels for each mutant were compared to wild-type (Table S1). Input RNA abundance was similar for all samples (Fig. 3B, input columns).

These IPs confirmed results with wild-type FBF-2 and Y479A, assayed previously by eCLIP (Carrick et al. 2024). Thus, wild-type FBF-2 and Y479A bound to all 17 known targets but not to the non-targets RNAs (Fig. 3B, IP WT and Y479A columns), and Y479A binding was weaker than wild-type for specific targets (e.g., *gld-1,* Fig. 3B, IP Y479A column; Fig. 3C). More importantly, the IPs demonstrated that TRM7^mut^ and TRM7^mut^ Y479A proteins did not bind RNA (Fig. 3B, IP TRM7^mut^ and TRM7^mut^ Y479A columns), much like the GFP negative control (Fig. 3B). Figure 3C quantitates results for binding to the *gld-1* target RNA and shows that abundance differences of immunoprecipitated RNA is significant. We conclude that the TRM7^mut^ destroys RNA binding *in vivo*.

### Conclusions and implications

This work investigates the *in vivo* significance of key RNA-binding residues in FBF-2, a paradigmatic PUF protein. Our results lead to three major conclusions. First, RNA-binding <u>is not</u> essential for one FBF-2–dependent biological function, maintenance of germline stem cells (GSCs): most TRM7^mut^ mutants maintain GSCs despite a lack of RNA binding. This unexpected result is important because it challenges the conventional wisdom that RBPs must bind to RNAs to exert biological activity. Second and in contrast to the first conclusion, RNA-binding <u>is</u> required for the sperm to oocyte cell fate switch. The differing first and second conclusions — RNA-binding required for one activity but not the other — highlight the likelihood of distinct mechanisms for the two major biological activities of a single PUF protein. Third, FBF-2 partner interactions are essential for GSC maintenance in TRM7^mut^ mutants, a conclusion that highlights the importance of PUF partner interactions and provides a clue about how an RNA-binding defective RBP may nonetheless retain biological function.

Figure 4 proposes a model to illustrate our thinking about how RNA recognition and protein partnerships may work together to accomplish *in vivo* FBF-2 functions. Central to this model is the idea that FBF-2 interacts with distinct Y479A-dependent partner complexes to achieve its two different biological functions. Figure 4A depicts wild-type FBF-2 binding to FBEs in its target RNAs as well as to Y479-dependent partners that modulate regulatory activity. The Y479-dependent partners chosen for illustration postulate one complex that represses RNAs in GSCs and a different complex that activates RNAs to promote the s/o switch. Although these examples are consistent with available evidence, other complexes and other factors may well contribute to these fate decisions. Figure 4B depicts TRM7^mut^ protein binding to partners but not to FBEs. We suggest that GSC-promoting partner complexes regulate RNAs without FBF-2 RNA binding (Fig. 4B, left), but that the switch-promoting partner complex requires FBF-2 RNA-binding (Fig. 4B, right). The challenge now is to understand how a GSC-promoting partner complex operates in the absence of FBF-2 RNA-binding. One simple model invokes involvement of a different RNA-binding protein that brings TRM7^mut^ protein to target RNAs (e.g. Qiu et al. 2024). Alternatively, an unknown RNA-independent role may be responsible. Figure 4C depicts the TRM7^mut^ Y479A double mutant protein, that no longer binds to either RNA or Y479-dependent partners and that no longer has biological activity. Our model thus proposes that FBF-2 operates via a multilayered regulatory mechanism, where different mechanisms compensate for one another, ensuring that essential biological processes are maintained even when one mode of interaction is compromised or downregulated. Such a multilayered mechanism may hold true for other PUF proteins and indeed other RBPs.

**Figure 4:**
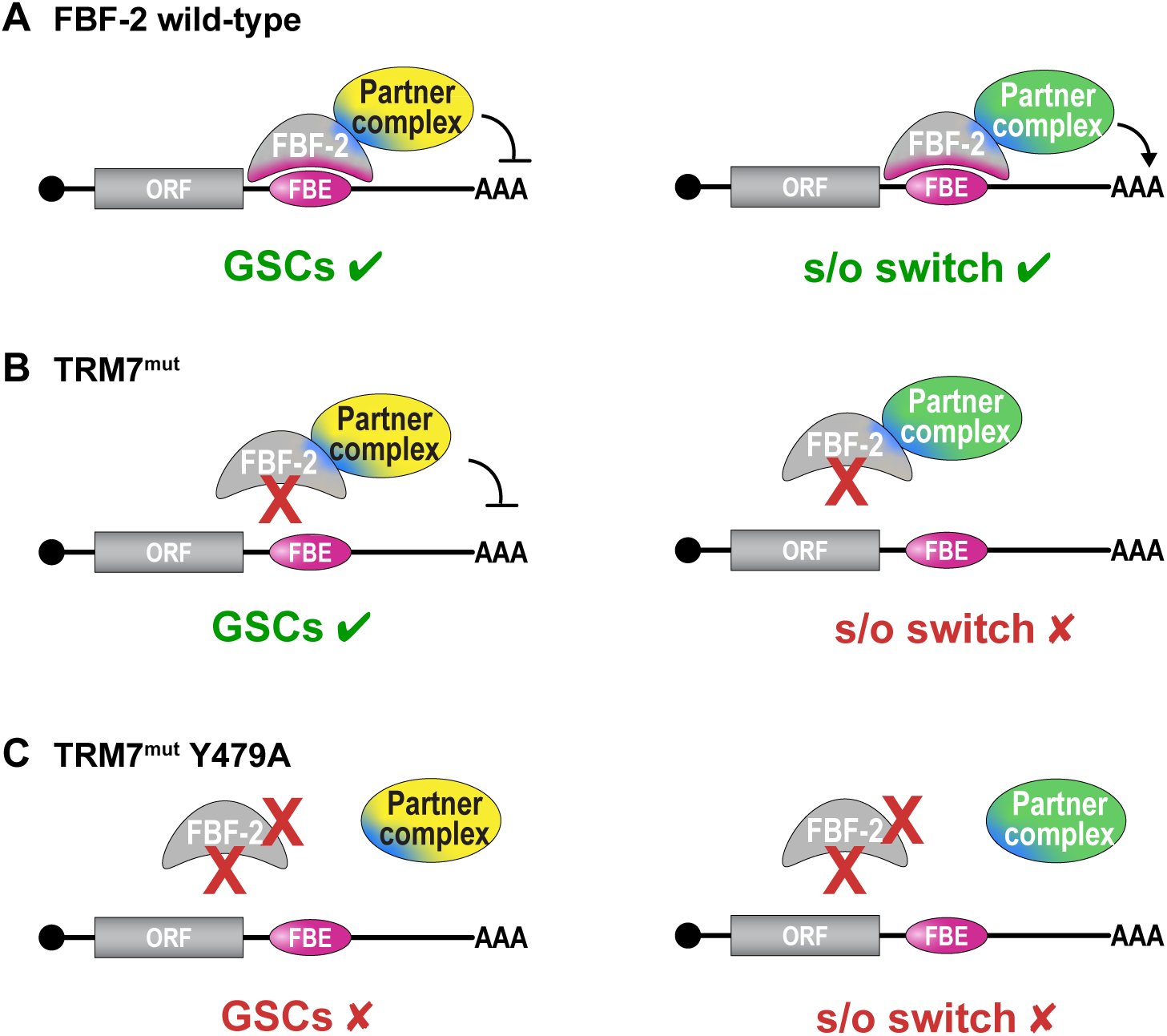
Models for RNA binding and protein partner effects on FBF-2 function. (A-C) Models to illustrate how RNA and protein partner interactions modulate FBF-2 function. Target RNAs (straight lines) with a cap (circle) at 5’ end, open reading frame (ORF) and 3’ untranslated region (3’ UTR) containing an FBF binding element (FBE). (A) Wild-type FBF-2 binds to an FBE in its target RNA and also interacts with protein partner complexes. Together these two binding interactions accomplish regulation sufficient for both GSC self-renewal (left) and the s/o switch fate (right). Both GSC-promoting (yellow) and s/o switch-promoting (green) partner complexes interact via the R7/R8 loop interface and Y479 (blue). (B) TRM7^mut^ does not bind FBE but does bind protein partner complexes. Partner complexes are sufficient to regulate RNA for GSC self-renewal (left) but not for the s/o switch (right). See text for further explanation. (C) TRM7^mut^ Y479A double mutant no longer binds RNA or protein partners, and thus loses its ability to regulate RNAs and accomplish either germline function.

PUF proteins, and RBPs more generally, are implicated in a wide spectrum of human diseases (Gennarino et al. 2015; Kapeli et al. 2017; Gennarino et al. 2018; Choi and Thomas-Tikhonenko 2021; Prashad and Gopal 2021). Disease-associated mutations can occur not only in RNA-binding residues, but also in residues located outside the RNA-binding domain, many within regions involved in protein-protein interactions. Our findings emphasize the critical need to understand how both RNA recognition and protein partnerships influence RBP function *in vivo*. While it might seem intuitive that a mutation disrupting RNA binding would eliminate an RBP’s biological activity, our work shows that this assumption is oversimplified. Consistent with that idea, RBPs with mutations in RNA binding residues can be oncogenic (Choi and Thomas-Tikhonenko 2021), a phenomenon that may rely on interactions with protein partners. A deeper understanding of RNA regulatory mechanisms is essential for unraveling the complexities of disease pathology and developing effective therapeutic strategies. Together our work highlights the importance of studying RBP function in the context of its binding partners in addition to its RNA-binding to target RNAs.

## Materials and Methods

### C. elegans maintenance

*Caenorhabditis elegans* were maintained by on NGM seeded with OP50 with standard techniques and grown at 20°C (Brenner 1974). Hermaphrodite animals were grown to 24 h past the L4 stage unless otherwise noted. Strains used are listed in table S2.

### CRIPSR-Cas9 mediated gene editing

New alleles were created by co-CRISPR editing using a CRISPR/Cas9 RNA-protein complex (Arribere et al. 2014; Paix et al. 2015; Dokshin et al. 2018). Animals were injected with a mix containing a gene-specific crRNA (5 μM, IDT-Alt-R), unc-58 crRNA (4 μM, IDT-Alt-R), tracRNA (4.5 μM, IDT), unc-58 repair oligo (1 μM, IDT), gene-specific repair oligo (5 μM, IDT) and Cas9 protein (3 μM, glycerol free, IDT). F1 progeny of injected hermaphrodites were screened for edits by PCR, homozygosed, sequenced and outcrossed against wild type prior to analysis. See table S3 for guide RNA and repair template sequences.

### mos1-mediated single-copy insertion (mosSCI)

DNA encoding *mex-5* promoter: eGFP with introns: 3xFLAG: *tbb-1* 3’UTR: *gpd-2* SL2 splice site: mCherry with introns: 3xmyc: PGL-1 RGG repeat: *tbb-1* intergenic region was cloned into pCFJ151 to create pJK1728. The transgene was inserted into the *ttTi5605* site on *LGII* of strain EG6699 using the *mos1-*mediated single copy insertion (mosSCI) method to generate *qSi100* (Frokjaer-Jensen et al. 2008). The presence of the transgene was verified by PCR and Sanger sequencing.

### Phenotypic analysis

Adult animals were scored as fertile or sterile using a dissecting scope. Sterile animals were then mounted on agarose pads and scored for germ cell morphology on a compound microscope. Progenitor zone length in germ cell diameters (gcd) was scored in DAPI-stained animals by counting germ cell diameters from the distal tip of the germline to the start of meiotic entry (Crittenden et al. 2023). Cells at the distal end of DAPI stained gonads were scored as GSCs, sperm, or variable. GSC germlines had a progenitor zone (PZ, appropriately sized cells followed by crescent shaped nuclei characteristic of early meiotic prophase). Sperm was identified by distinctive highly condensed DNA. ’Variable’ gonads contained enlarged nuclei, crescents or ambiguous cells at the distal end.

### RIP-qPCR

Strains JK5081, JK5810, JK5984, JK6593, and JK6737 were cultivated at 20°C and grown to early adulthood (24 h after L4) in all RIP-qPCR replicates. Developmental stage was evaluated with a Leica Wild M3Z stereoscope to score body size and stage-specific marks (e.g., vulva formation). Animals were kept on standard NGM plates and fed *E. coli* OP50 as previously described (Stiernagle 2006). Age-synchronized first stage larvae (L1) were obtained by bleach synchronizing gravid adults by standard methods (Lewis and Fleming 1995). Briefly, gravid adults were treated with 2:1 bleach:4N NaOH to isolate embryos. Embryos were resuspended in M9 buffer (per 1L of buffer: 6 g Na_2_HPO_4_, 3 g KH_2_PO_4_, 5 g NaCl, 1 ml of 1 M MgSO_4_) without food in a ventilated Erlenmeyer flask at 20°C for 20 h. L1s were pelleted at 2500 rcf for 2 min, washed twice with 15 ml of M9, and distributed to 10 cm NGM plates pre-equilibrated to 20°C. Plates were pre-seeded with 1.5 ml of 40x concentrated OP50. Three biological replicates of each genotype were obtained. At least 100,000 animals were used per replicate, and each plate contained no more than 10,000 worms per plate.

Once animals reached L4 + 24-h stage, live worms were quickly rinsed from plates into a 15 ml falcon tube with cold M9 + 0.01% Tween-20 (M9Tw), washed once with cold M9Tw, pelleted at 200 RCF in cold M9Tw, and transferred by glass pipet to a 2 ml tube, and snap frozen in liquid nitrogen. Pellets were stored at -80°C.

Pellets were thawed by adding 800 μL ice-cold lysis buffer (50 mM HEPES pH 7.5, 100 mM NaCl, 1% NP-40, 0.1% SDS, 0.5% sodium deoxycholate, 1x Roche cOmplete, EDTA-free protease inhibitor cocktail, 1 U/μL SUPERase•In RNase inhibitor) and rocking for 20 min at 4°C. Thawed pellets were centrifuged at 1000 RCF at 4°C for 1 min and washed three times with 800 μL cold lysis buffer. One ml of lysis buffer was added to the pellet along with a 5-mm stainless steel ball (Retsch). Lysis was performed at 4°C with a Retsch 400 MM mill mixer (3x 10-min cycles of 30 Hz). Cracking of tube lid was prevented by adding 2 small pieces of duct tape to the lid just prior to lysis.

Complete tissue lysis was confirmed by observing a small aliquot of lysate at 40x magnification. Lysate was clarified at 16,000 RCF for 15 min at 4°C. Protein concentration was determined using Bio-Rad Protein Assay Dye (Bio-Rad #5000006) and measuring absorbance at 595 nm on a Bio-Rad SmartSpec 3000.

To prepare antibody conjugated beads, 10 μg mouse αFLAG was incubated with 4.5 mg protein G Dynabeads (Novex, Life Technologies, #10003D) for 60 min at RT. Beads were then washed 2x with lysis buffer. 20 mg of total protein was incubated with the antibody-bead mixture for 4h at 4°C. Beads were washed three times with lysis buffer, and then three times with wash buffer (same as lysis but with 500 mM NaCl). Successful IP was confirmed by analyzing 1% of elution by Western blot. 1% of beads were resuspended in (2% (w/v) SDS, 0.1% βME, 10% glycerol, 50 mM Tris pH 8) and incubated for 10 min at 100°C and analyzed by SDS-PAGE (4-20% acrylamide gel). For primary antibodies, blots were incubated overnight at 4°C at the following dilutions: αFLAG M2 (1:1000; Sigma-Aldrich, Cat# F1804), αGAPDH (1:10,000; Proteintech, Cat# 60004-1-Ig). For secondary antibody, blots were incubated for 1 hour at RT with HRP-conjugated anti-mouse (1:10,000, Jackson ImmunoResearch, Cat# 115-035-003).

RNA was purified by adding 500 μL acid-phenol:chloroform:isoamyl alcohol (125:24:1, Invitrogen AM9722, PCA) to the remaining beads (still in last wash). Samples were mixed by gentle shaking and were separated by centrifugation for 15 min at 15,000 r.p.m. at 4°C. The aqueous layer was removed (∼500μL) and further extracted by three additional extractions (1x PCA followed by 2x chloroform:isoamyl alcohol). After the extractions, the aqueous layer was removed and ∼1 mL of 100% ethanol was added to the samples, which were gently mixed and incubated at −50°C for >1 h. RNA was pelleted by centrifugation for 30 min at 15,000 r.p.m. at 4°C. Pellets were washed once with ∼70% ethanol and resuspended in 43 μL of water. 8 units of TURBO DNase (Life Technologies #AM2238) was then added for 1 h at 37°C. RNA was purified using the GeneJet RNA Purification kit (Thermo Fisher Scientific #K0732) and eluted in 30 μL of water. RNA samples were stored at −80°C until use.

RNA was converted to cDNA with SuperScript III First-Strand Synthesis System (Invitrogen #18080051) using random hexamers as primers. Quantitative PCR was carried out in technical triplicate in a Roche Lightcycler 480 using the LightCycler 480 SYBR Green I Master (Roche #04887352001). Average C_T_ of the technical replicates for each biological replicate is given in table S1. Primers used for each gene tested are listed in table S4. Comparative C_T_ method (2^-ΔΔCT^) was used to calculate relative amounts of RNA present using *rps-25* to normalize and making all comparisons to wild-type (Schmittgen and Livak 2008). Significance was calculated in GraphPad Prism 10.0.0 using one-way ANOVA and Dunnett’s multiple comparisons test (all compared to wild-type).

### Immunostaining and imaging

Animals were staged at mid-L4 and grown for 24 h at 20°C and then processed for immunostaining. We immunostained gonads as described with minor modifications (Crittenden et al. 2023). Gonads were dissected in PBS containing 0.1% (v/v) Tween-20 (PBST) and 0.25 mM levamisole. Gonads were fixed in 4% (w/v) paraformaldehyde in PBST for 10 min, then permeabilized in 0.2% (v/v) Triton-X in PBST. Next, gonads were incubated for at least 30 min in blocking solution (30% goat serum in PBST), washed 3 times with PBST, and incubated overnight at 4° with primary antibodies diluted in blocking solution. After washing, secondary antibodies were diluted in blocking solution and incubated with samples for at least 1 h. To visualize DNA, DAPI was included with the secondary antibody at a final concentration of 1 ng/μl. After washing, samples were mounted in ProLong Gold (#P36930; Thermo Fisher Scientific) and cured overnight to several days before imaging. All steps were performed at room temperature unless otherwise indicated. Antibody concentrations were as follows: αFLAG M2 (1:1000; Sigma-Aldrich, Cat# F1804), αSP56 (1:100; Sam Ward (Ward et al. 1986)), αPH3 (1:1000, Cell Signaling Technology Cat #9706), αMouse-Alexa647 (1:1000; Molecular Probes/Invitrogen Cat# A-31571), αRabbit-Alexa488 (1:1000; Molecular Probes/Invitrogen Cat# A-21206). Imaging was performed on a Leica SP8 confocal microscope.

### Competing Interest Statement

The authors declare no competing interests.

## Acknowledgements

The authors thank members of the Kimble and Wickens labs for insightful discussions throughout the course of this work. We thank Jane Selegue, Jadwiga Forster, and Peggy Kroll-Conner for technical assistance and Scott Aoki for constructing JK5081. We thank Laura Vanderploeg for assistance with figure preparation. Some strains were provided by the CGC, which is funded by the NIH Office of Research Infrastructure Programs (P40 OD010440). This work was supported by the National Science Foundation Graduate Research Fellowship Program under grant numbers DGE-1256259 and DGE-1747503 to B.H.C., NIH R01 GM50942 to M.W., and NIH R01 GM134119 to J.K. Any opinions, findings, and conclusions or recommendations expressed in this material and those of the authors do not necessarily reflect the views of the National Science Foundation.

## Author Contributions

Conceptualization, B.H.C; formal analysis, B.H.C., S.L.C. and M.L.; investigation, B.H.C., S.L.C., M.L., S.J.C.D.S. and J.K.; resources, B.H.C. and M.L.; data curation, B.H.C. and S.L.C.; writing – original draft, B.H.C. and J.K.; writing – review & editing, B.H.C., S.L.C, M.W. and J.K.; funding acquisition, B.H.C., M.W. and J.K.

---

**Table S2.** – Strains used in this study

| Strains | Genotype | Simplified nomenclature | Reference |
| --- | --- | --- | --- |
| N2 | Wild type | Wild-type | Brenner 1974 |
| JK3022 | *fbf-1(ok91) II* | *fbf-1(∅)* | Crittenden et al. 2002 |
| JK3101 | *fbf-2(q738) II* | *fbf-2(∅)* | Lamont et al. 2004 |
| JK3107 | *fbf-1(ok91) fbf-2(q704) / mIn1[mls14 dpy-10(e128)] II* | *fbf-1(∅) fbf-2(∅)* | Crittenden et al. 2002 |
| JK5081 | *qSi100 III; unc-119(ed3)* | eGFP<sup>3xFLAG</sup>, PGL-1 RGG repeat<sup>3xmyc</sup> | This study |
| JK5810 | *fbf-2(q945) II* | FBF-2<sup>3xFLAG</sup> | Ferdous et al. 2023 |
| JK5984 | *fbf-2(q1011) II* | FBF-2(Y479A)<sup>3xFLAG</sup> | Carrick et al. 2024 |
| JK5986 | *fbf-1(ok91) fbf-2(q1023) / mIn1[mls14 dpy-10(e128)] II* | *fbf-1(∅) FBF-2(Y479A)<sup>3xFLAG</sup>* | Carrick et al. 2024 |
| JK6578 | *fbf-1(ok91) fbf-2(q1261) / mIn1[mls14 dpy-10(e128)] II* | *fbf-1(∅) FBF-2(S453A, H454A, E457A)<sup>3xFLAG</sup> aka TRM7<sup>mut</sup>* | This study |
| JK6593 | *fbf-2(q1262) II* | FBF-2(S453A, H454A, E457A)<sup>3xFLAG</sup> aka TRM7<sup>mut</sup> | This study |
| JK6596 | *fbf-1(ok91) fbf-2(q1272) / mIn1[mls14 dpy-10(e128)] II* | *fbf-1(∅) FBF-2(S453A, H454A, E457A, Y479A)<sup>3xFLAG</sup> aka TRM7<sup>mut</sup> Y479A* | This study |
| JK6658 | *fbf-1(ok91) fbf-2(q1285) / mIn1[mls14 dpy-10(e128)] II* | *fbf-1(∅) FBF-2(N415A, Y416A, Q419A, S453A, H454A, E457A)<sup>3xFLAG</sup> aka TRM6<sup>mut</sup> TRM7<sup>mut</sup>* | This study |
| JK6737 | *fbf-2(q1295) II* | FBF-2(S453A, H454A, E457A, Y479A)<sup>3xFLAG</sup> aka TRM7<sup>mut</sup> Y479A | This study |

**Table S3.**
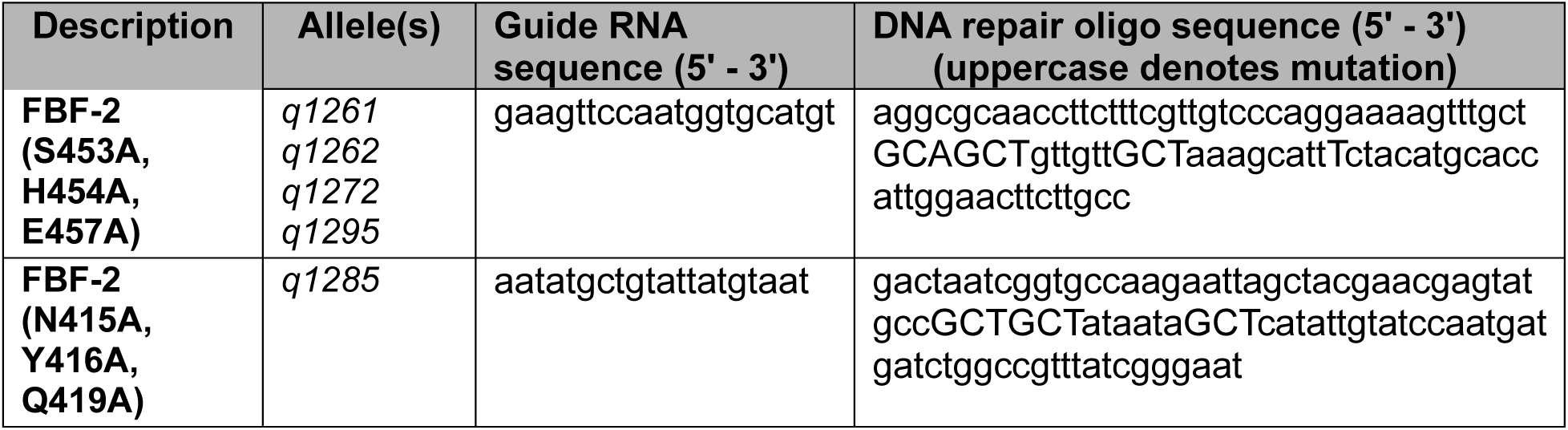
– CRISPR-Cas9 gene editing reagents

**Table S4.**
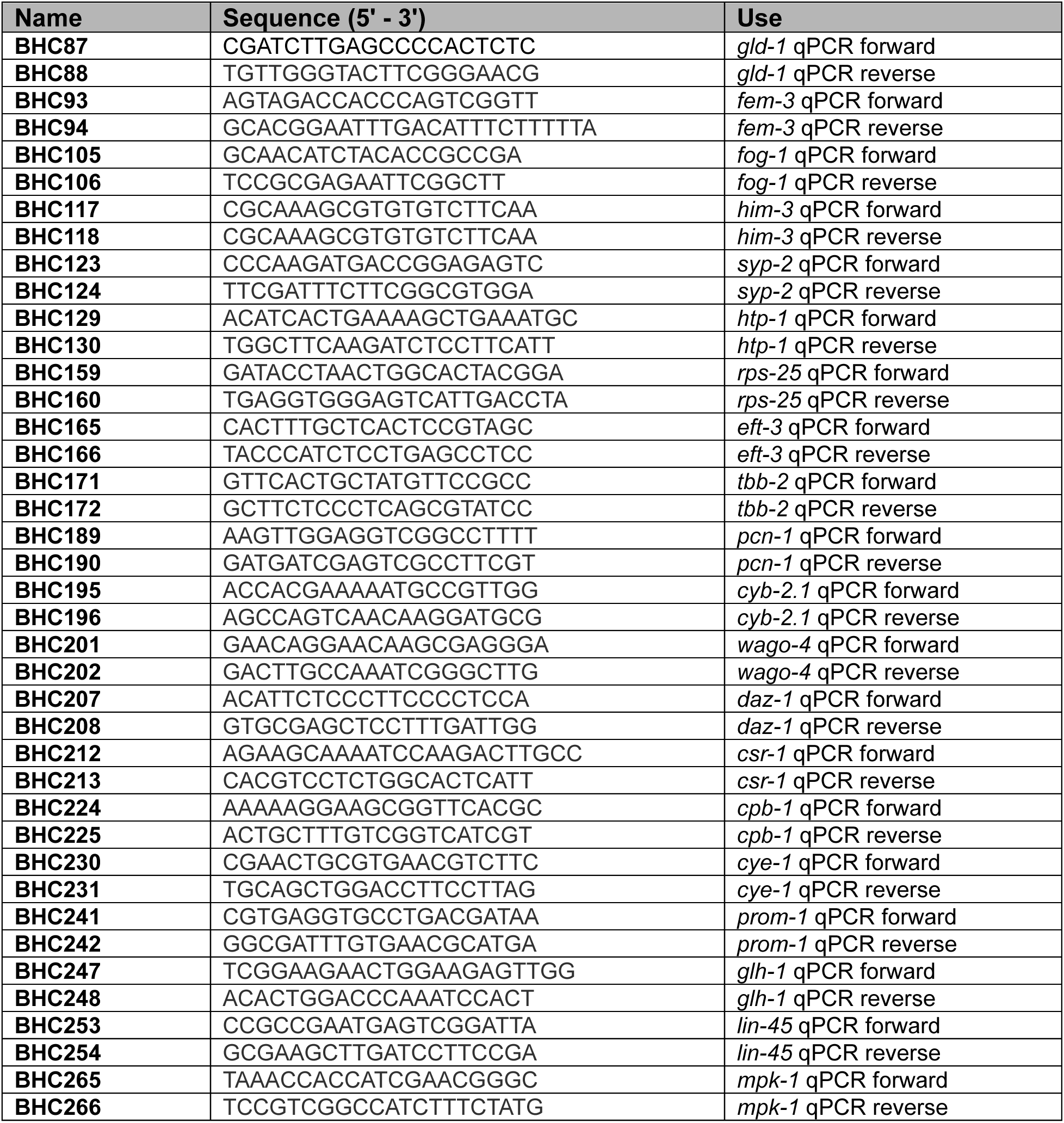
– qPCR primers

## References

1. Arribere JA, Bell RT, Fu BX, Artiles KL, Hartman PS, Fire AZ. 2014. Efficient marker-free recovery of custom genetic modifications with CRISPR/Cas9 in Caenorhabditis elegans. Genetics 198: 837–846.

2. Brenner S. 1974. The genetics of Caenorhabditis elegans. Genetics 77: 71–94.

3. Campbell ZT, Bhimsaria D, Valley CT, Rodriguez-Martinez JA, Menichelli E, Williamson JR, Ansari AZ, Wickens M. 2012a. Cooperativity in RNA-protein interactions: global analysis of RNA binding specificity. Cell Rep 1: 570–581.

4. Campbell ZT, Menichelli E, Friend K, Wu J, Kimble J, Williamson JR, Wickens M. 2012b. Identification of a conserved interface between PUF and CPEB proteins. J Biol Chem 287: 18854–18862.

5. Carrick BH, Crittenden SL, Chen F, Linsley M, Woodworth J, Kroll-Conner P, Ferdous AS, Keles S, Wickens M, Kimble J. 2024. PUF partner interactions at a conserved interface shape the RNA-binding landscape and cell fate in Caenorhabditis elegans. Dev Cell 59: 661–675 e667.

6. Choi PS, Thomas-Tikhonenko A. 2021. RNA-binding proteins of COSMIC importance in cancer. J Clin Invest 131.

7. Corley M, Burns MC, Yeo GW. 2020. How RNA-Binding Proteins Interact with RNA: Molecules and Mechanisms. Mol Cell 78: 9–29.

8. Crittenden SL, Bernstein DS, Bachorik JL, Thompson BE, Gallegos M, Petcherski AG, Moulder G, Barstead R, Wickens M, Kimble J. 2002. A conserved RNA-binding protein controls germline stem cells in Caenorhabditis elegans. Nature 417: 660–663.

9. Crittenden SL, Seidel HS, Kimble J. 2023. Analysis of the C. elegans Germline Stem Cell Pool. Methods Mol Biol 2677: 1–36.

10. Daigle JG, Lanson NA, Jr., Smith RB, Casci I, Maltare A, Monaghan J, Nichols CD, Kryndushkin D, Shewmaker F, Pandey UB. 2013. RNA-binding ability of FUS regulates neurodegeneration, cytoplasmic mislocalization and incorporation into stress granules associated with FUS carrying ALS-linked mutations. Hum Mol Genet 22: 1193–1205.

11. Dokshin GA, Ghanta KS, Piscopo KM, Mello CC. 2018. Robust Genome Editing with Short Single-Stranded and Long, Partially Single-Stranded DNA Donors in Caenorhabditis elegans. Genetics 210: 781–787.

12. Edwards TA, Pyle SE, Wharton RP, Aggarwal AK. 2001. Structure of Pumilio reveals similarity between RNA and peptide binding motifs. Cell 105: 281–289.

13. Friend K, Campbell ZT, Cooke A, Kroll-Conner P, Wickens MP, Kimble J. 2012. A conserved PUF-Ago-eEF1A complex attenuates translation elongation. Nat Struct Mol Biol 19: 176–183.

14. Frokjaer-Jensen C, Davis MW, Hopkins CE, Newman BJ, Thummel JM, Olesen SP, Grunnet M, Jorgensen EM. 2008. Single-copy insertion of transgenes in Caenorhabditis elegans. Nat Genet 40: 1375–1383.

15. Gennarino VA, Palmer EE, McDonell LM, Wang L, Adamski CJ, Koire A, See L, Chen CA, Schaaf CP, Rosenfeld JA et al. 2018. A Mild PUM1 Mutation Is Associated with Adult-Onset Ataxia, whereas Haploinsufficiency Causes Developmental Delay and Seizures. Cell 172: 924–936 e911.

16. Gennarino VA, Singh RK, White JJ, De Maio A, Han K, Kim JY, Jafar-Nejad P, di Ronza A, Kang H, Sayegh LS et al. 2015. Pumilio1 haploinsufficiency leads to SCA1-like neurodegeneration by increasing wild-type Ataxin1 levels. Cell 160: 1087–1098.

17. Gerstberger S, Hafner M, Tuschl T. 2014. A census of human RNA-binding proteins. Nat Rev Genet 15: 829–845.

18. Goldstrohm AC, Seay DJ, Hook BA, Wickens M. 2007. PUF protein-mediated deadenylation is catalyzed by Ccr4p. J Biol Chem 282: 109–114.

19. Kapeli K, Martinez FJ, Yeo GW. 2017. Genetic mutations in RNA-binding proteins and their roles in ALS. Hum Genet 136: 1193–1214.

20. Lamont LB, Crittenden SL, Bernstein D, Wickens M, Kimble J. 2004. FBF-1 and FBF-2 regulate the size of the mitotic region in the C. elegans germline. Dev Cell 7: 697–707.

21. Lewis JA, Fleming JT. 1995. Basic culture methods. Methods Cell Biol 48: 3–29.

22. Lunde BM, Moore C, Varani G. 2007. RNA-binding proteins: modular design for efficient function. Nat Rev Mol Cell Biol 8: 479–490.

23. Paix A, Folkmann A, Rasoloson D, Seydoux G. 2015. High Efficiency, Homology-Directed Genome Editing in Caenorhabditis elegans Using CRISPR-Cas9 Ribonucleoprotein Complexes. Genetics 201: 47–54.

24. Prashad S, Gopal PP. 2021. RNA-binding proteins in neurological development and disease. RNA Biol 18: 972–987.

25. Qiu C, Bhat VD, Rajeev S, Zhang C, Lasley AE, Wine RN, Campbell ZT, Hall TMT. 2019. A crystal structure of a collaborative RNA regulatory complex reveals mechanisms to refine target specificity. Elife 8.

26. Qiu C, Crittenden SL, Carrick BH, Dillard LB, Costa Dos Santos SJ, Dandey VP, Dutcher RC, Viverette EG, Wine RN, Woodworth J et al. 2024. A higher order PUF complex is central to regulation of C. elegans germline stem cells. *bioRxiv*.

27. Qiu C, Zhang Z, Wine RN, Campbell ZT, Zhang J, Hall TMT. 2023. Intra-and inter-molecular regulation by intrinsically-disordered regions governs PUF protein RNA binding. Nat Commun 14: 7323.

28. Schmittgen TD, Livak KJ. 2008. Analyzing real-time PCR data by the comparative C(T) method. Nat Protoc 3: 1101–1108.

29. Siomi H, Choi M, Siomi MC, Nussbaum RL, Dreyfuss G. 1994. Essential role for KH domains in RNA binding: impaired RNA binding by a mutation in the KH domain of FMR1 that causes fragile X syndrome. Cell 77: 33–39.

30. Stiernagle T. 2006. Maintenance of C. elegans. WormBook: 1–11.

31. Suh N, Crittenden SL, Goldstrohm A, Hook B, Thompson B, Wickens M, Kimble J. 2009. FBF and its dual control of gld-1 expression in the Caenorhabditis elegans germline. Genetics 181: 1249–1260.

32. Valley CT, Porter DF, Qiu C, Campbell ZT, Hall TM, Wickens M. 2012. Patterns and plasticity in RNA-protein interactions enable recruitment of multiple proteins through a single site. Proc Natl Acad Sci U S A 109: 6054–6059.

33. Wang X, McLachlan J, Zamore PD, Hall TM. 2002. Modular recognition of RNA by a human pumilio-homology domain. Cell 110: 501–512.

34. Wang X, Zamore PD, Hall TM. 2001. Crystal structure of a Pumilio homology domain. Mol Cell 7: 855–865.

35. Ward S, Roberts TM, Strome S, Pavalko FM, Hogan E. 1986. Monoclonal antibodies that recognize a polypeptide antigenic determinant shared by multiple Caenorhabditis elegans sperm-specific proteins. J Cell Biol 102: 1778–1786.

36. Webster MW, Stowell JA, Passmore LA. 2019. RNA-binding proteins distinguish between similar sequence motifs to promote targeted deadenylation by Ccr4-Not. Elife 8.

37. Weidmann CA, Goldstrohm AC. 2012. Drosophila Pumilio protein contains multiple autonomous repression domains that regulate mRNAs independently of Nanos and brain tumor. Mol Cell Biol 32: 527–540.

38. Weidmann CA, Qiu C, Arvola RM, Lou TF, Killingsworth J, Campbell ZT, Tanaka Hall TM, Goldstrohm AC. 2016. Drosophila Nanos acts as a molecular clamp that modulates the RNA-binding and repression activities of Pumilio. Elife 5.

39. Wu J, Campbell ZT, Menichelli E, Wickens M, Williamson JR. 2013. A protein.protein interaction platform involved in recruitment of GLD-3 to the FBF.fem-3 mRNA complex. J Mol Biol 425: 738–754.

40. Zamore PD, Williamson JR, Lehmann R. 1997. The Pumilio protein binds RNA through a conserved domain that defines a new class of RNA-binding proteins. RNA 3: 1421–1433.

41. Zhang B, Gallegos M, Puoti A, Durkin E, Fields S, Kimble J, Wickens MP. 1997. A conserved RNA-binding protein that regulates sexual fates in the C. elegans hermaphrodite germ line. Nature 390: 477–484.

